# The Human Pangenome’s sequence conservation reveals a landscape of polymorphic structural variations

**DOI:** 10.1101/2022.10.06.511239

**Authors:** HoJoon Lee, Stephanie U. Greer, Dmitri S. Pavlichin, Christopher R. Hughes, Bo Zhou, Tsachy Weissman, Human Pangenome Reference Consortium, Hanlee P. Ji

**Author notes:** A complete list of authors and contributions appears at the end of the paper. To whom correspondence should be addressed. Hanlee P. Ji, Division of Oncology, Department of Medicine – Stanford University School of Medicine, CCSR 1115, 269 Campus Drive, Stanford, CA 94305-5151.

## Abstract

The Human Pangenome is a new reference build that addresses many of the limitations of the current reference. The first release is based on 94 high-quality haploid assemblies from individuals with diverse backgrounds. To facilitate the Pangenomes adoption in the wider research community, we conducted a multiple genome comparative analysis against the current GRCh38 reference. We applied a k-mer indexing strategy to identify highly conserved sequences that are omnipresent across all Pangenome assemblies, the reference and CHM13, a reference assembly with telomere-to-telomere chromosomes. Pan-conserved tag segments provided an informative set of universally conserved sequences. Examining the intervals between pairs of these segments defined highly conserved segments of the genome versus ones that have structurally related polymorphisms. We identified a Pangenome landscape of 60,764 polymorphic intervals with unique and geo-ethnic features. Overall, this study of the Pangenome revealed the conserved versus divergent features including the landscape of polymorphic structural variants.

## INTRODUCTION

The human genome reference has been instrumental in the discovering the genetic basis of human diseases and an essential component for a wide variety of genetic and genomic applications. However, the current reference **(GRCh38)**has limitations^1^. Specifically, it lacks haploid features, has gaps in the sequence and provides limited representation of genetic diversity across different human populations. These limitations make it more difficult to characterize specific, complex genome features relevant to human disease such as structural variations **(SVs)**^2,3^. Recently, the telomere-to-telomere **(T2T)**consortium produced a complete assembly from the haploid cell line CHM13hTERT **(CHM13)**^4^. On an expanded scale, Human Pangenome Reference Consortium **(HPRC)**is constructing a new reference based on hundreds of high accuracy haploid assemblies, representing the whole genome sequences of multiple individuals with broad genetic diversity^5^. The initial HPRC Pangenome release includes 94 haploid assemblies^6^. This new reference eliminate gaps, incorporates complex genomic sequence features, and capture a greater breadth of human genome diversity. The new features are based on dramatic improvements in sequencing technology and assembly construction. As a result, the HPRC Pangenome provides a high-quality, complete representation of human genomes and enables identification of a greater breadth of variants compared to GRCh38.

To facilitate the adoption of the Human Pangenome among the genetics research community, one must define the different properties of the new versus the current reference. Since the Pangenome is comprised of multiple haploid assembles, sequence alignment to GRCh38 involves making multiple comparisons across different genomes. This approach requires significant computing resources and poses numerous challenges^7^. As an efficient and straightforward solution, we developed an indexing strategy which identifies highly conserved short sequences, referred to as k-mers, across different assemblies^8^. K-mers - short sequences with length “k”, typically in range of tens of bases - have many advantages for genome comparisons that include rapid encoding, scalability of processing multiple genome sequences and annotation of sequence features from diverse sets of assemblies. K-mer indexing enabled us to identify a series of pan-conserved segments, which were identical among all individuals contributing to the Pangenome and we evaluated the lengths between pan-conserved segments to deterime genomic regions where structural variation was present and polymorphic among the Pangenome. Thus, we identified the Pangenome’s structurally conserved versus divergent sequence features.

## RESULTS

### Pan-conserved segments among the HPRC assemblies and references

The HPRC released 47 phased and diploid assemblies from four superpopulations in addition to one Ashkenazim Jewish **(Supplementary Table 1)**: 24 African **(AFR)**, 16 Admixed American **(AMR)**, 5 East Asian **(ESA)**, and 1 South Asian **(SAS)**^6^. There were 28 females and 19 males. The number of contigs per a given haploid genome ranged from 236 to 817 with an average of 408 per haploid assembly and the majority of contigs were longer than 100 kb. The HPRC achieved a consistent sequencing coverage across various sequencing platforms and this high-quality sequence data provided a high-quality input for genome assemblies^6^. We focused on the autosomes and excluded the X and Y chromosomes due to their absence among a large subset of the genomes that were analyzed - both maternal and paternal haploid assemblies from 28 females lacked chromosome Y while all paternal haploids from 19 males lacked chromosome X.

We compared the 94 HPRC assemblies with the GRCh38 reference genome and the CHM13 - a gapless telomere-to-telomere assembly using our k-mer indexing method^8^. For indexing all haploid genomes, the input assembly sequence was parsed into its constituent 31-mers using a sliding window and associated with their locations and frequencies in the input assembly **(Methods)**. These short sequences can be efficiently compared across multiple genome data sets as we previously demonstrated^9^. We used 31-mers in this study because of multiple advantages as we previously described^10^ including that the vast majority (82.8%) of 31-mers from GRCh38 were unique within an edit distance of 2 bases **(Supplementary Fig. 1)**.

Among the 96 genomes and the two references, we identified a total of ~1.62 x 10^9^ 31-mers **(Supplementary File 1)** that have the following properties: (1) they occur only once in each haploid genome; (2) they are present with the same uniqueness feature across all of the genomes **(Methods)**. Based on the GRCh38 coordinates, we observed that 1.55 x 10^9^ (98.7%) of the identified 31-mers had consecutive positions **(Supplementary Fig. 2A)**. Consecutive overlapping 31-mers form longer segments of sequence that were present across all haploid assemblies - we refer to these extended sequences as pan-conserved segment tags **(PSTs) (Fig. 1A)**. The set of PSTs were distributed across all chromosomes **(Fig. 1B)** and specific genome features including exonic, intronic and intergenic regions **(Fig. 1C)**. As expected, centromeres and the acrocentric regions in the p arms of specific chromosomes had a lower density of PSTs - this was a result of their highly repetitive sequence structure. As we noted previously, we focused on the autosomes given the sex differences among the contributors. We confirmed that PSTs were present in all haploid genomes **(Supplementary Note)**.

**Figure 1.**
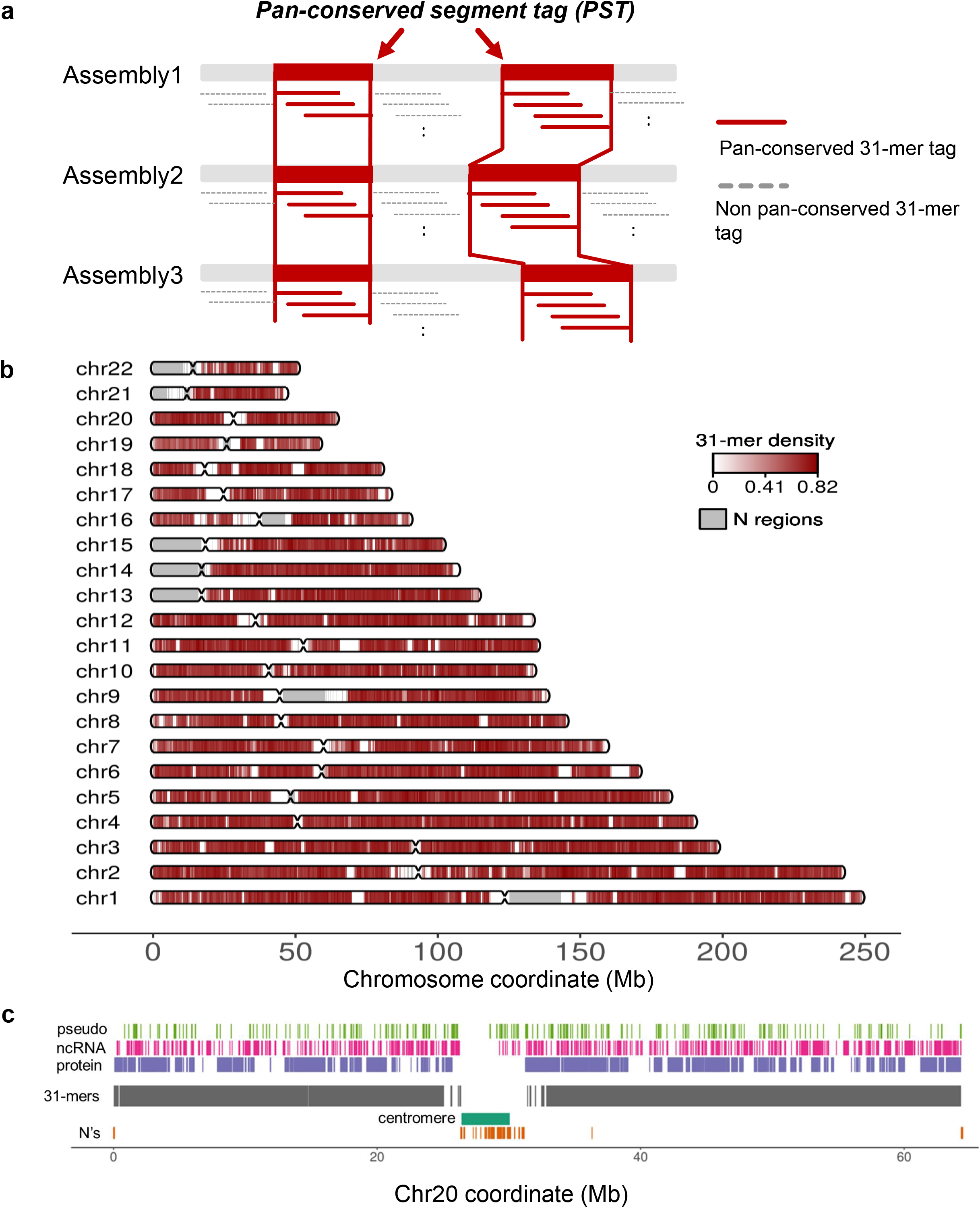
Identification of pan-conserved segment tag in HPRC assemblies and their properties based on GRCh38 coordinates. (a) We define PST as when the set of consecutive unique sequence is present in all assemblies. (b) The distribution of PSTs on GRCh38. The density of PSTs was calculated in 500 kb window; number of pan-conserved 31-mers/size of window. (c) The distribution of PSTs across the different types of genomics regions on chr20. Annotate genomic regions with N’s as N regions.

### Characteristics of the pan-conserved segment tags (PSTs)

The median length of an individual PST was 65 bp with ranging from 31 bp (from an individual 31-mer) up to a maximum size of 5.26 kilobase **(kb) (Supplementary Fig. 2B)**. There were 24 PSTs with length greater than 2.5 kb and mapped to genome coordinates in both GRCh38 and CHM13 **(Supplementary Table 2)**. The longest PST appeared on chromosome 5q31.3 where two genes (*PURA* and *IGIP*) with single exons are located. Thus, this same segment was the same among all assemblies. In addition, we confirmed that this 5q31.3 segment lacked any variants with a population frequency of 1% or higher according to Genome Aggregation Database **(gnomAD)**^11^. This result is an indicator of sequence conservation among humans. Furthermore, the sequence alignment of this region from 30 different species of mammals showed that this segment was highly conserved across all primates, mice, and dogs^12,13^. The other 24 longer PST had similar characteristics of high sequence conservation. Based on the coordinates of CHM13, we observed similar PST characteristics which included: (1) the density of pan-conserved 31-mers tag, (2) distance between tandem pan-conserved 31-mers tag, and (3) the length distributions of PSTs **(Supplementary Fig. 3)**.

### Intervals between PST pairs among the Pangenome assemblies

The interval length between tandem cis-based pairs of PSTs provided a way to determine the presence of structural variation **(SV)** among the individual haploid assemblies. Systematically examining all PST pairs, we determined the interval lengths among all haploid genomes and determined the differences in interval lengths compared to the reference. Variation in interval length for any given haploid genome are indicators of SVs **(Fig. 2A)**. A constant interval length across haploid genomes implies the absence of SVs while a structural variation introduces changes in the interval length.

**Figure 2.**
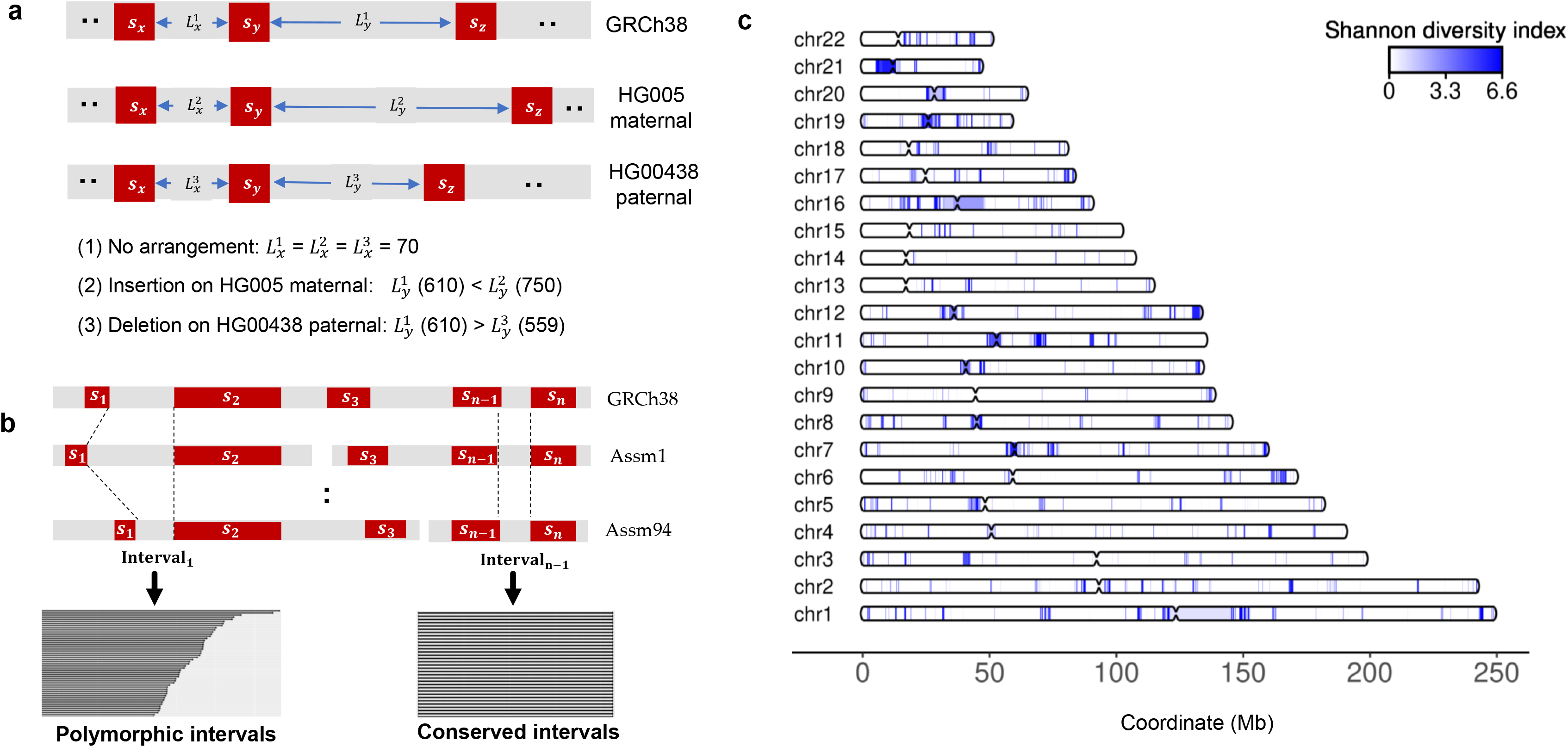
Intervals between pan-conserved sequences tags (PSTs). (A) The three types of interval lengths relative to the interval length on GRCh38: (1) No arrangement: interval length on an assembly is identical to the length on GRCh38, (2) Insertion: interval length on an assembly is larger than the length on GRCh38, and (3) Deletion: interval length on an assembly is less than the length on GRCh38, (B) Measuring length of intervals between adjacent pan-conserved sequence pair after sorting them by GRCh38 coordinates. A small number (< 0.00001%) of tandem pairs of PST were on different contigs for a given haploid genome thanks to high quality of HPRC assemblies (*i.e.* S_2_ and S_3_ in Assm1), (D) The distribution of polymorphic intervals with Shannon diversity index of the divergent lengths across assemblies. S_i_ indicates the i^th^ PST while Assm stands for assembly.

This evaluation involved the two steps **(Fig. 2B)**: (1) all PSTs were sorted for each chromosome based on the GRCh38 coordinates using a p to q arm orientation and (2) we calculated the length of the interval sequence between any two tandem PST pairs within a given assembly contig. We conducted this process for all 94 HPRC haploid assemblies and constructed a data matrix where the columns represented a given haploid genome and the rows represented the lengths between consecutive PSTs **(Supplementary Fig. 4)**. This matrix contained approximately 13.5 million rows with the interval lengths between tandem pairs of PSTs for a given haploid assembly **(Supplementary File 2)**. This matrix provided a rapid way to identify intervals with the same versus different lengths across 94 assemblies.

PSTs have limits when mapped to the contigs with either the p and q telomeres. Only limited number of contigs for a given haploid genome had this type of PST. Similarly, the Pangenome has not released complete chromosome assemblies - the breaks between contigs prevented some tandem segments from being compared (**Fig. 2B**). This represented only a small fraction of the HPRC assemblies.

### Conserved intervals across the Pangenome

Among the 13.5 million intervals, we identified approximately 11.3 million (83.6%) where the sequence lengths were identical among all 94 assemblies **(Supplementary File 3)**. The uniform interval length implied that the intervening sequences were the same, albeit there could be variants such as SNPs which do not change the number of bps. These ‘conserved intervals’ indicate the segments of the Pangenome with a high degree of structural conservation. We used the Matched Annotation from NCBI and EMBL-EBI **(MANE)**resource to determine coding regions for each conserved interval^14^. Approximately 51 Mb (67.5%) of exonic regions and 905 Mb (73 %) of genic regions including introns overlapped with the conserved intervals. Five hundred and twenty conserved regions were longer than 10 Kb **(Supplementary Table 3)**. The longest region (chr13:102729323-102751307) spanned 22 Kb across the gene *CCDC168*. More than half of the conserved regions occurred outside of genic regions and the longest one spanned 18 Kb at chr2:63078147-63096212 where there was no reported protein-encoding gene^13^.

### Divergent interval lengths point to loci with polymorphic structural variations

We identified 60,763 (0.45%) intervals that had divergent lengths of 50 bp or greater compared to the GRCh38 for at least one haploid assembly among all assemblies **(Supplementary Table 4)**. In general, longer interval lengths had higher levels of divergence **(Supplementary Note)**. The median divergent lengths in the size range of 100 bp to 1 kb bracket was the highest frequency category (47.2%) of polymorphic interval **(Supplementary Table 5)**. As we show per our results, these divergent lengths defined the location of structural polymorphisms that were present throughout the Pangenome.

We observed that 46,516 polymorphic intervals overlapped with repeats per a comparison with the RepeatMasker annotation (4.0.6). There were 12,906 intervals located within repeat sequences and motifs. For the size range up to 100kb, the LINE/L1s had the most frequent overlap. For the size range of 100 kb to 1 Mb and 1 Mb to 10 Mb, SINE/Alus and LTR/ERVL-MaLRs had the most frequent ovlerap. Interestingly, we observed different trends for shorter divergent lengths; simple repeat for 50 to 100 bp, SINE/Alu for 100 to 500 bp and LINE/L1 for 500 to 1000 bp. Additional categories are listed in **Supplementary Table 6**.

Sixty one polymorphic intervals were located within the coding exon regions, and thereby changed the lengths of the coding sequences in 39 genes, such as, *MUC6, MYH8* and *ATG9B*. The vast majority (>95%) of these different interval lengths were related to in-frame variants **(Supplementary Table 7)**. Citing an example, all 94 assemblies had different lengths for the exon 31 of *MUC6* compared to GRCh38.

Next, we characterized the polymorphic intervals based on the frequency of the divergent lengths among the haploid genomes: (1) singletons which are present in only one haploid assembly, (2) low frequency intervals (1-5%), and (3) high frequency intervals (>5%) **(Supplementary Table 5)**. Interestingly, singletons were very frequent (28.7%). There was one class of singletons that were directly related to the GRCh38. We identified 253 intervals which had identical lengths among all 94 assemblies but differed only for GRCh38 - this category were indicators of a reference limitation. Citing an example, all 94 assemblies had an additional 252 bp in the last exon of *ZNF676* on chr19 - only the GRCh38 reference lacked this feature. The remaining singletons were indicators of potential SVs that were private to an individual haploid genome. In general, the number of total polymorphic intervals and singletons per assembly was higher among the AFR genome compared to other populations **(Supplementary Table 8)**. This observation is consistent with what has been reported by other studies^15^.

We measured the extent of variability for these polymorphic intervals by (1) the Shannon-Wiener diversity index and (2) an inter-quartile range **(IQR)**. The diversity index provides a quantitative metric regarding the extent of different interval lengths while the IQR value provided information on the magnitude of length variation **(Methods)**. The median of Shannon-Wiener diversity index for intervals was 0.42. The values ranged from 0 (when all 94 assemblies have an interval length) to 6.56 (when all 94 assemblies have different lengths) **(Fig. 2D)**. Notably, three intervals had different lengths for all 93 individual assemblies. The first one was located at chr11:11246448-11247212 with median different lengths of 4.4 Kb overlapped with LTR and simple repeats. The second one occurred at chr13:111793323-111843451 with median different lengths of 94 Kb did not overlap with any genes or repeats. Within the interval of chr15:34278120-34586890 with median different lengths of 5.2 Kb, there were several genes including SLC12A6 and NOP1 and repeats such as SINE, LINE, and simple repeats. As an example, we showed all divergent interval lengths for 94 assemblies on chr18 in **Figure 3A**. The other chromosomes are shown in **Supplementary Fig 6**. We highlighted three examples: (1) the most polymorphic interval on chr18 **(Fig. 3B)**, (2) an interval with largest divergent length **(Fig. 3C)**, and (3) an interval length with a high frequency **(Fig. 3D)**.

**Figure 3.**
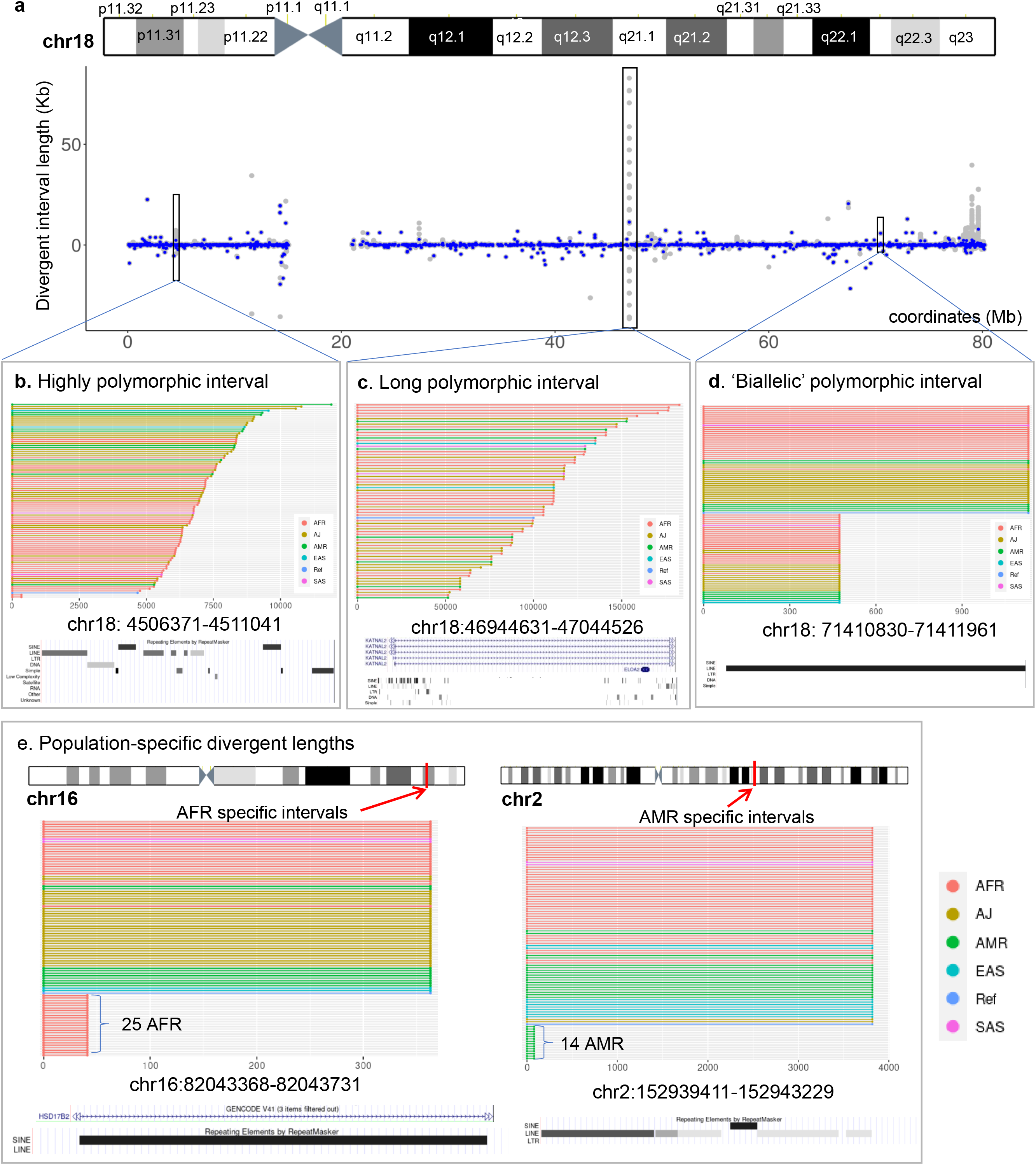
Polymorphic intervals on chromosome 18. (A) The locations of polymorphic intervals across chromosomes18. Blue dots indicate the median of interval lengths while gray dots indicate the interval length of an assembly, (B) Highly polymorphic interval of 4.67 kb size based on GRCh38 had 92 different lengths with a diversity index of 6.51, (C) Long polymorphic interval had the highest IQR of divergent length relative to the reference interval size of 47 kb. (D) A biallelic polymorphic interval with high frequency (0.457) has a binomial distribution of different lengths (658 bp deletion for 43 assemblies; no changes for 51 assemblies). The entire region of this interval is annotated as LINE by RepeatMasker. (E) Population-specific intervals with divergent lengths only for AFR and AMR. The interval at chr16:82043368-82043731 had a deletion of 322 bp on intron 1 of SD17B2. This deletion was present exclusively among the 25 AFR assemblies where 12 of them were homozygous for 6 individuals.

Finally, we identified some of the divergent intervals associated with specific biogeographic populations **(Supplementary Table 9)**. We found 381 divergent intervals with polymorphic lengths present in only a single super-population **(Fig. 3E)**. We cite examples in which the divergent interval lengths were present in 10 or more assemblies: 376 were specific to the AFR superpopulation; 5 were specific to the AMR super-population. We also observed 59 intervals with the reverse attribute. For example, the chr12:102848406-102848539 interval had a 70bp deletion among 59 assemblies, but this deletion was not present among all 10 EAS assemblies.

### Polymorphic intervals compared to reported SVs

We examined whether the polymorphic intervals were overlapping with SVs reported from 1000 Genomes Project **(1KGP)**^16^. We observed that 13,624 (93.8%) out of 14,526 simple SVs from 1KGP^17^ overlapped with one of polymorphic intervals **(Table 1)**. We excluded insertions from analysis since this class of SVs are the most challenging to accurately call and still vastly underrepresented in gold standard call sets^18,19^. The high overlapping indicates that the analysis of the Pangenome provided a way to identify regions of the genome that contain SV polymorphisms in the population.

**Table 1.** The number of 1000 Genome Project SVs overlapping with the polymorphic intervals. We compared the polymorphic intervals with 22,288 simple SVs with >1% frequency from 1KGP. Simple SVs include deletion (DEL), duplication (DUP), insertion (INS), and inversion (INV). The polymorphic intervals were grouped by their median of different lengths. The numbers in bold indicate the most frequent length difference in each SV type.

| **Interval lengths difference** | **DEL (11965)** | **DUP (2368)** | **INV (193)** |
| --- | --- | --- | --- |
| <100bp | 1887 | 689 | 42 |
| 100 bp – 1 Kb | 6063 | 980 | 82 |
| 1 Kb – 10 Kb | 2540 | 301 | 24 |
| 10 Kb – 100 Kb | 398 | 102 | 6 |
| 100 Kb– 1 Mb | 190 | 39 | 6 |
| 1 Mb – 10 Mb | 81 | 34 | 1 |
| 10 Mb – 100 Mb | 131 | 23 | 5 |
| Total | 11290 (94.4%) | 2168 (91.6%) | 166 (86%) |

### Benchmarking divergent intervals as indicators of structural variants

To demonstrate that the polymorphic intervals were indicators of SVs, we examined the HG002 genome for presence of deletions, insertions and other SVs. HG002 is part of the Pangenome and this individual has also undergone an extensive genomic analysis by the Genome in a Bottle **(GIAB)** Consortium. Per the GIAB analysis, HG002 had 250 SVs located in the vicinity of medically relevant genes^20^. We compared 220 SVs that had a size of 50 bp or greater with our polymorphic intervals. Remarkably, all 220 SVs were located within the polymorphic intervals found in the Pangenome **(Supplementary Table 10)**. The size of most SVs (74.5%) had the same lengths as those described by divergent interval lengths. Another subset of SVs (11%) had minor difference in lengths by less than 15 bp. For a small subset of the SVs, the reported size from the GIAB benchmark did not match our divergent interval length. These were 31 intervals that had a median length of 100kb - these long intervals contained multiple SV structures that led to a difference.

### Visualizing the structure of SVs using constituent sequences from the Pangenome

Providing an example, we using the constituent 31-mers of SVs to visualize the structure of different classes of SVs including insertions, deletions, duplications, inversion and more complex rearrangements. This process involved using a simple dot matrix plot with the two axis representing the GRCh38 and the specific haploid assembly. We plotted the position of the 31-mers that spanned the divergent interval **(Fig. 4A)**. We showed six different SV classes occurring in these divergent intervals **(Fig. 4B–4G)**. As expected, longer intervals indicated either insertions or tandem duplication while shorter intervals indicated either a deletion or a complex SV. Furthermore, we were able to characterize the structure of a highly complex SV identified in CHM13. This complex SV involved nine different structural change that included multiple insertions, deletions and inversions **(Fig. 5)**.

**Figure 4.**
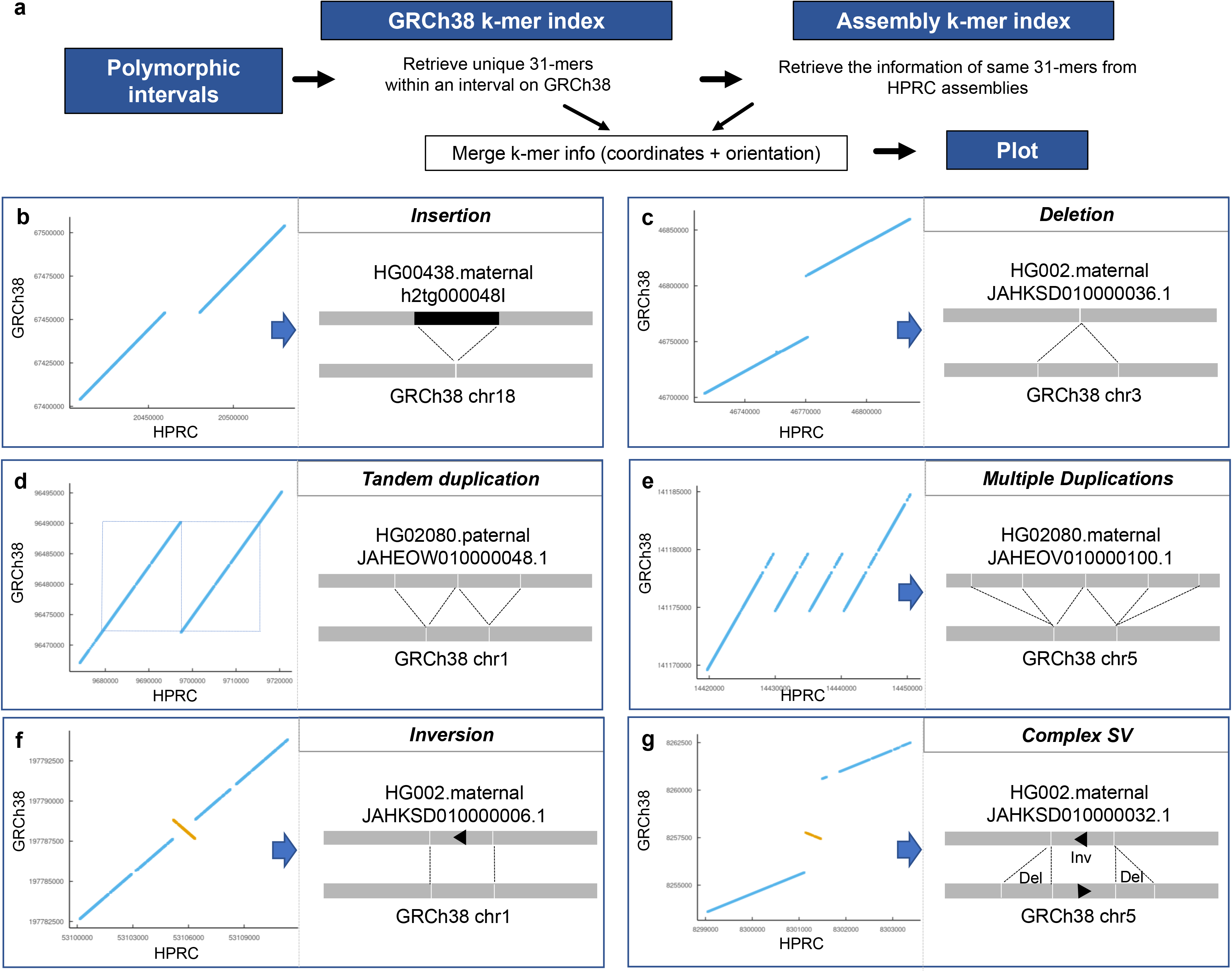
The SV plots using 31-mers from polymorphic intervals. (a) The scheme of plotting 31-mers from both reference and query assemblies to depict SVs. Examples of different types of SVs are shown in: (b) insertion, (c) deletion, (d) tandem duplication, (e) multiple duplications, (f) inversion, and (g) complex SV.

**Figure 5.**
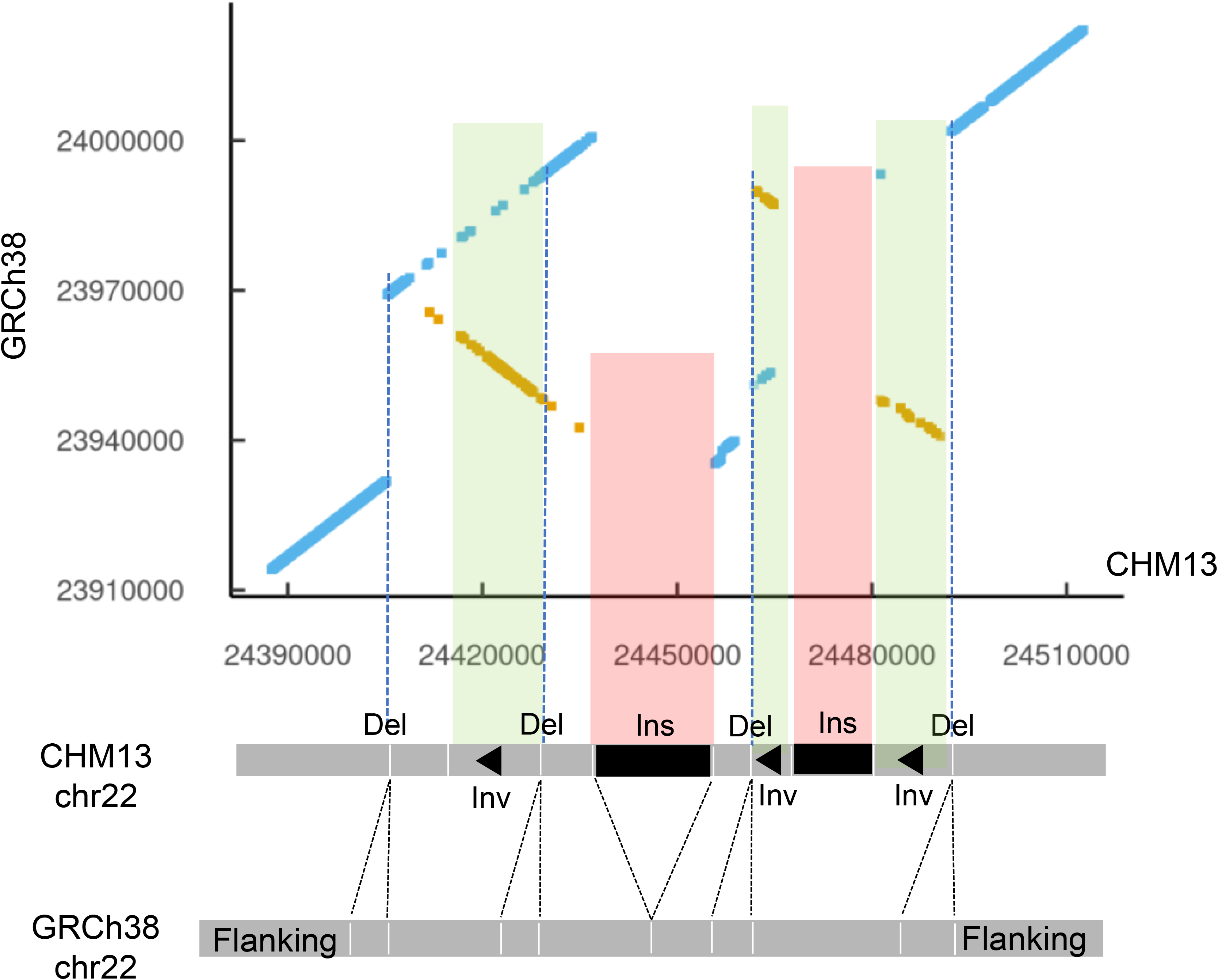
Anatomy of complex structural variant (SV). We demonstrate that an SV plot displays the various components within a complex SV.

## DISCUSSION

The Human Pangenome Reference Consortium has released its first draft of human pangenome derived from 94 haploid assemblies of 47 individuals with diverse genetic background^6^. The Pangenome new features provide increased representation of genomic and geographic diversity and haplotype structure. With its improved sequence assembly, it addresses the limitations of current linear reference genome, GRCh38. To promote the adoption of pangenome reference, it is critical to make comparison these assemblies with the reference genomes. Thus, there is a need for new approaches to enable the research community to utilize pangenome reference in sequencing analysis.

As we have described, this k-mer indexing approach enables one to conduct multi-genome comparisons efficiently and in a highly scalable fashion. For identifying conserved versus divergent sequence features, this approach has several advantages over conventional sequence alignment, particularly in a multiple genome comparisons. First, PSTs are independent of any individual genome’s coordinates - this allows one to use the coordinates for a given assembly or any other reference such as GRCh38. Demonstrating this feature, we have provided all our results in CHM13 coordinates as supplementary files in addition to ones based on GRCh38. Another advantage is that it can be used on incomplete assemblies and long-read sequences. This feature also allows direct comparison among different genomes. For instance, we observed an average of 14,522 deviated lengths relative to GRCh38 per a haploid assembly while we observed an average of 11,936 divergent interval lengths between maternal and paternal haploids from an individual.

Divergent interval lengths based on PSTs points to potential structural variants such as deletions. As noted among our results, these divergent lengths revealed a set of genome loci that are highly polymorphic. Beyond the identification of polymorphic loci in the genome, this approach and related resource can be used to rapidly visualize SV structure. For example, segmental duplications look like insertions that results in longer intervals in this study. In addition, the divergent interval lengths indicates the presence of potential SVs. Juxtaposition of k-mers involving SVs from two assemblies can reveal the general sturcture SVs including complex ones **(Fig. 4 and 5)**. Specifically, one can use other classes of 31-mers without some of the stringent criteria metrics and apply this expanded set for identifying rearrangement features. For example, these 31-mers with looser sequence characteristics can distinguish tandem duplications with insertions. PSTs with their unique segments also provide an accessible way of visualizing complex SVs. For example, S_1_S’_4_S’_3_S’_2_S_5_ describes the inversion of S_2_S_3_S_4_ and S_1_S_2_S_3_S_4_S_1_S_5_ describes interspersed duplication of S_1_ when a reference looks like S_1_S_2_S_3_S_4_S_5_ where S_1_, S_2_, S_3_, S_4_, S_5_ represent PST/unique segments. Negative interval lengths may pinpoint the rearranged PST due to SVs including inversion. For future studies, we will develop computational tools to determine the breakpoints of SVs with their k-mer plots and PSTs.

As a resource for the research community, we provide our interval matrix of 94 HPRC assemblies against GRCh38/CHM13 in a BED format. This file is readily accessible and is formatted such that it can used across a variety of different applications. Other researchers can take advantage of this matrix of Pangenome conserved/divergent sequences to see which regions contain their variants of interest. Furthermore, our method using PSTs to split *de novo* haploid assemblies in same manner enables systematic characterization of genomic conservation and divergence.

The HPRC will be expanding the Pangenome reference to include more haploid assemblies^5^. This approach is readily scalable across hundreds of genomes. Thus, we can readily update this resource for the final release. In summary, the comparison of available haploid assemblies relative to the reference genomes in a timely manner will enable using the Pangenome resource and holds the potential to further accelerate new genetic discoveries.

## Methods

### The sequences of human genome assemblies

In this study, we analyzed a total of 47 individuals with 94 haploid human genome assemblies in addition to two references from the following sources; 1) GRCh38, 2) CHM13, and 3) 94 HPRC haploid assemblies. We obtained the following assemblies from the National Center for Biotechnology Information **(NCBI)**. From Genbank, we downloaded the 94 haploid assemblies from 47 individuals, which generated by HPRC (year 1 freeze Genbank). The accession number of assemblies are in **Supplementary Table 1**. The HPRC samples underwent whole genome sequencing that included long sequence reads (Pacific Biosciences, Oxford Nanopore), optical mapping (Bionano) and high coverage short read sequencing. The consortium developed a bioinformatic pipeline with multiple quality control metrics. The production process included evaluating the completeness, contiguity, base-level quality, and phasing accuracy of each haploid assembly.

### The structural variant data

For benchmarking HG002, we downloaded GIAB CMRG benchmark containing medically relevant SVs. In addition, we downloaded 22,288 SVs (deletions, duplications, insertions, and inversions) called from high-coverage whole-genome sequencing of the expanded 1000 Genomes Project cohort. We overlapped them with the polymorphic intervals using bedtools (v2.27): bedtools intersect -wa -wb -b $polymorphic_intervals.bed -a $sv_1kgp.bed. In instances where a SV overlaps with multiple polymorphic intervals, we selected polymorphic intervals with the maximum base-pair overlap.

### K-mer indexing of assemblies

To characterize these assemblies, we indexed them using our k-mer-based indexing strategy^8^. A given assembly are indexed in two steps **(Supplementary Fig. 7A)**.

Step 1: retrieve the sequence of k-mers with sliding window with 1 bp increments.

Step 2: Associate the k-mers with genomics positions and their frequencies.

Citing an example, for the first substring of length 3 at position 1 is AAT and second substring at position 2 is ATA. We repeat this process from first position to (*n-k+1*)^th^ position where n = the length of assembly and k is the length of substring. We counted canonical k-mers, where k-mers are identical to their reverse complement, and selected a sequence based on lexicographical order. For instance, CGA is selected for S_6_ instead of TCG. The frequency of AAT is 2 since AAT appears at position 1 and position 4 while the frequency of ATA is 1.

We repeat this process with all assemblies and build a large index for all assemblies of interest (**Supplementary Fig. 7B**). This index enables us to retrieve the information through sequences from indexed assemblies such as the list of assemblies with query sequences and their locations in each assembly.

### Defining the properties of k-mers in a haploid genome assembly

For this study, we defined three categories of k-mer metrics derived from a given haploid genome assembly; (1) total set of k-mers, (2) the non-duplicated, distinct k-mers, (3) unique k-mers (**Supplementary Fig. 7A**). Given an assembly A of length L, the first metric describes the total number of k-mers of length k that are present in a given genome assembly (A). Thus, this is a count for every sequence substring. We denote the sequence substring of length k at position i in A as *S*_*i*_.

**Total set of k-mers** = {*S*_1_, *S*_2_…, *S*_*L* – *k* + 1_}

The second metric involves identify and counting all of the different sequences from the total number of k-mers which are not duplicated (distinct). We denote i^th^ k-mers as *k*_*i*_ after sorting all substrings in lexicographic order.

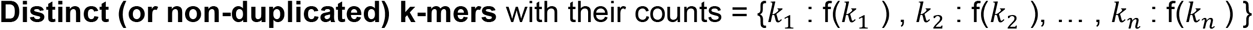

where f(*k*_*i*_) is the count of observed *k*_*i*_ in A.

The next k-mer metrics involves the sequence substring of length k which are present only once (unique) from the total set of k-mers for a given haploid assembly. Therefore, unique k-mers in A are all *k*_*i*_ with f(*k*_*i*_) = 1

**Unique k-mers**; *Uniq*_*A*_ = {*all k*: f(*k*_*i*_) = 1}.

### The definition of “pan-conserved k-mers tag”

From multiple haploid assemblies from different individuals, we define a highly conserved subset of k-mers that have the following properties: (1) are non-duplicated and thus distinct per a given haploid genome; (2) found only once per a given haploid genome and thus are unique; (3) are observed across all of the assemblies with the same properties. For the last point, this k-mer subset represents an intersection across all assemblies. This k-mers from this intersection are conserved across all individual genomes that were included in this total set of assemblies.

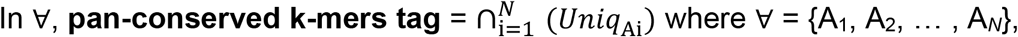

Namely, the k-mers which are not-duplicated, unique per a given haploid assembly and have these same properties across all individuals in a collection of genomes. To facilitate referring to this set of k-mers, we will use an acronym that describes these properties: pan-conserved sequences tag (PST). This definition has advantages in that it can be adjusted for new haploid genome assemblies as they become available.

As an example in **Supplementary Fig. 7B**, we will consider an example from Assembly 1. The sequence AATAATCGA has total 7 substrings of length 3; {*S*_1_, *S*_2_ .., *S*_7_} and 4 distinct 3-mers since the sequences of *S*_1_ and *S*_4_ as well as *S*_6_ and *S*_7_ are identical. There are 3 unique 3-mers out of 5 distinct 3-mers, *Uniq*_A1_ = {*ATA*, *ATC*, *TAA*}. We repeated same process with Assembly 2 and 3 to identify *Uniq*_A2_ = {*ATC*, *CAA, CAC, GAC*} and *Uniq*_A3_ = {*AAT*, *ATC*, *CGA*, *GAA*}. We identify pan-conserved 3-mers tag by *Uniq*_A1_ ∩ *Uniq*_A2_ ∩ *Uniq*_A3_ = {*ATC*}. Among 3 assemblies, *ATC* is only pan-conserved tag sequence since it is unique in assembly 1, 2, and 3 respectively. *AAT* is not pan-conserved 3-mers tag since *ATT* ∉ *Uniq*_A1_. All other unique 3-mers are not present in all assemblies. For instance, *CAA* is not pan-conserved 3-mer tag because *CAA* ∊ *Uniq*_A2_, but *CAA* ∉ *Uniq*_A1_ and *CAA* ∉ *Uniq*_A3_.

### The matrix of interval lengths between the PSTs across 94 assemblies

The genomic arrangements were measured by the length of intervals between adjacent PST. Possible genomic rearrangements in a given assembly can be estimated based on the difference in the interval length (deviated lengths) as follows:

- Deletion: 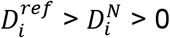
- Insertion: 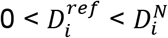

Calculation of all interval lengths involves the following steps: First, we sorted PST by their coordinates based on a reference, which is GRCh38 in this study. Second, we retrieve the location of PST on contigs from an assembly. Third, we measured the lengths between adjacent PST, *S*_*i*_ and *S*_*i*+1_, for i = 1 to n-1 for an assembly. The lengths between *S*_*i*_ and *S*_*i*+1_ is measured by their starting positions. The distance cannot be measured when *S*_*i*_ and *S*_*i*+1_ are on different contigs.

Since all contigs from an assembly are not in the same orientation, we determined the orientation of contigs based on the majority of signs of length. For instance, if there are more minus lengths than positive ones, we switched all signs of lengths.

### Measuring the variability of interval lengths between adjacent PSTs across 94 assemblies

We identified PSTs by combining consecutive pan-conserved 31-mers tag into a segment. The n^th^ interval was defined from last 31-mers in n^th^ PST and first 31-mers in n+1^th^ PST. The PST with less than 50 pan-conserved 31-mers tag were represented only by their first pan-conserved 31-mers tag. Each interval has the lengths for 94 HPRC assemblies. The extent of variability for polymorphic-intervals was measured by (1) Shannon-Wiener diversity index and (2) inter-quartile range **(IQR)**. First, we used a Python library called “scikit-bio”, which defines Shannon-Wiener diversity index as 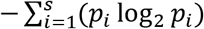 where s is the number of different distinct lengths and *p*_*i*_ is the proportion of the lengths of i. The Shannon-Wiener diversity shows how many different lengths are in an interval across 94 HPRC assemblies. Second, the inter-quartile range is calculated by Python NumPy library as follows: IQR = Q3 - Q1 where Q3 is 3^rd^ quartile value and Q1 is 1^st^ quartile value. The IQR value provides information on the magnitude of deviated lengths.

## DATA AVAILABILITY

Two human genome references used in this study are GCA_000001405.15_GRCh38_no_alt_analysis_set.fna for GRCh38 and chm13.draft_v1.1.fasta for CHM13.

All HPRC haploid assemblies are available at https://github.com/human-pangenomics/HPP_Year1_Assemblies. GIAB CMRG benchmark containing medically relevant SVs are available at: https://ftp-trace.ncbi.nlm.nih.gov/ReferenceSamples/giab/release/AshkenazimTrio/HG002_NA24385_son/CMRG_v1.00/GRCh38/StructuralVariant/.

The SVs from 1KGP (1KGP_3202.Illumina_ensemble_callset.freeze_V1.vcf.gz) are obtained from http://ftp.1000genomes.ebi.ac.uk/vol1/ftp/data_collections/1000G_2504_high_coverage/working/20210124.SV_Illumina_Integration/

The sequences and coordinates of pan-conserved sequences tag (PSTs) are publicly available at https://dna-discovery.stanford.edu/publicmaterial/datasets/pangenome/. In addition, all 13.5 M of measured intervals lengths across all assemblies were available at the same site.

## SOFTWARE AND CODE AVAILABILITY

The web version of KmerKeys is available at the following URL: https://kmerkeys.dgi-stanford.org/. The R scripts used in this study are available at the following GitHub: https://github.com/compbio/pan-conserved_segments.

## FUNDING

All authors were supported by National Institutes of Health grant [U01HG01096]. HPJ received additional support from the Clayville Foundation.

## ACKNOWLEDGEMENT

We would like to thank Billy Lau, Xiangqi Bai and Shubham Chandak for helpful discussions. We also thank Alison Almeda and Jung Yoo for providing comments on the manuscript.

## AUTHOR INFORMATION

Authors and Affiliations

**Division of Oncology, Department of Medicine, Stanford University School of Medicine, Stanford, CA, 94305, United States**

HoJoon Lee, Stephanie U. Greer, Dmitri S. Pavlichin, Hanlee P. Ji

**Department of Psychiatry and Behavioral Sciences, Stanford University School of Medicine, Stanford, CA 94305, USA**

Bo Zhou & Christopher R. Hughes

**Department of Genetics, Stanford University School of Medicine, Stanford, CA 94305, USA**

Christopher R. Hughes & Bo Zhou

**Department of Electrical Engineering, Stanford University, Palo Alto, CA, 94304, United States**

Tsachy Weissman & Hanlee P. Ji

**Consortia**

Human Pangenome Reference Consortium

Wen-Wei Liao^1,2,3,*^, Mobin Asri^4,*^, Jana Ebler^5,*^, Daniel Doerr^5^, Marina Haukness^4^, Glenn Hickey^4^, Shuangjia Lu^3^, Julian K. Lucas^4^, Jean Monlong^4^, Haley J. Abel^6^, Silvia Buonaiuto^7^, Xian H. Chang^4^, Haoyu Cheng^8,9^, Justin Chu^8^, Vincenza Colonna^7,10^, Jordan M. Eizenga^4^, Xiaowen Feng^8,9^, Christian Fischer^10^, Robert S. Fulton^1^, Shilpa Garg^11^, Cristian Groza^12^, Andrea Guarracino^13^, William T Harvey^14^, Simon Heumos^15,16^, Kerstin Howe^17^, Miten Jain^18^, Tsung-Yu Lu^19^, Charles Markello^4^, Fergal J. Martin^20^, Matthew W. Mitchell^21^, Katherine M. Munson^14^, Moses Njagi Mwaniki^22^, Adam M. Novak^4^, Hugh E. Olsen^4^, Trevor Pesout^4^, David Porubsky^14^, Pjotr Prins^10^, Jonas A. Sibbesen^23^, Chad Tomlinson^1^, Flavia Villani^10^, Mitchell R. Vollger^14,24^, Lucinda L Antonacci-Fulton^1^, Gunjan Baid^34^, Carl A. Baker^14^, Anastasiya Belyaeva^34^, Konstantinos Billis^20^, Andrew Carroll^34^, Pi-Chuan Chang^34^, Sarah Cody^1^, Daniel E. Cook^34^, Omar E. Cornejo^35^, Mark Diekhans^4^, Peter Ebert^5^, Susan Fairley^20^, Olivier Fedrigo^36^, Adam L. Felsenfeld^37^, Giulio Formenti^36^, Adam Frankish^20^, Yan Gao^38^, Carlos Garcia Giron^20^, Richard E. Green^39,40^, Leanne Haggerty^20^, Kendra Hoekzema^14^, Thibaut Hourlier^20^, Hanlee P. Ji^41^, Alexey Kolesnikov^34^, Jan O. Korbel^42^, Jennifer Kordosky^14^, HoJoon Lee^41^, Alexandra P. Lewis^14^, Hugo Magalhães^5^, Santiago Marco-Sola^43,44^, Pierre Marijon^5^, Jennifer McDaniel^29^, Jacquelyn Mountcastle^36^, Maria Nattestad^34^, Nathan D. Olson^29^, Daniela Puiu^45^, Allison A Regier^1^, Arang Rhie^28^, Samuel Sacco^46^, Ashley D. Sanders^47^, Valerie A. Schneider^48^, Baergen I. Schultz^37^, Kishwar Shafin^34^, Jouni Sirén^4^, Michael W. Smith^37^, Heidi J. Sofia^37^, Ahmad N. Abou Tayoun^49,50^, Françoise Thibaud-Nissen^48^, Francesca Floriana Tricomi^20^, Justin Wagner^29^, Jonathan M. D. Wood^17^, Aleksey V. Zimin^45,51^, Alice B. Popejoy^52^, Guillaume Bourque^25,26,27^, Mark JP Chaisson^19^, Paul Flicek^20^, Adam M. Phillippy^28^, Justin M. Zook^29^, Evan E. Eichler^14,30^, David Haussler^4,30^, Erich D. Jarvis^31,30^, Karen H. Miga^4^, Ting Wang^32^, Erik Garrison^10,+^, Tobias Marschall^5,+^, Ira Hall^3,33,+^, Heng Li^8,9,+^, Benedict Paten^4,+^

1 McDonnell Genome Institute, Washington University School of Medicine, St. Louis, MO 63108, USA

2 Department of Medicine, Washington University School of Medicine, St. Louis, MO 63110, USA

3 Department of Genetics, Yale University School of Medicine, New Haven, CT 06510, USA

4 UC Santa Cruz Genomics Institute, University of California, Santa Cruz, 1156 High St, Santa Cruz, CA, USA

5 Institute for Medical Biometry and Bioinformatics, Medical Faculty, Heinrich Heine University Düsseldorf, Düsseldorf, Germany

6 Division of Oncology, Department of Internal Medicine, Washington University School of Medicine, St. Louis, MO 63110, USA

7 Institute of Genetics and Biophysics, National Research Council, Naples 80111, Italy

8 Department of Data Sciences, Dana-Farber Cancer Institute, Boston, MA 02215, USA

9 Department of Biomedical Informatics, Harvard Medical School, Boston, MA 02215, USA

10 Department of Genetics, Genomics and Informatics, University of Tennessee Health Science Center, Memphis, TN 38163, USA

11 Department of Biology, University of Copenhagen, Denmark

12 Quantitative Life Sciences, McGill University, Montreal, Québec H3A 0C7, Canada

13 Genomics Research Centre, Human Technopole, Milan 20157, Italy

14 Department of Genome Sciences, University of Washington School of Medicine, Seattle, WA 98195, USA

15 Quantitative Biology Center (QBiC), University of Tübingen, Tübingen 72076, Germany

16 Biomedical Data Science, Department of Computer Science, University of Tübingen, Tübingen 72076, Germany

17 Tree of Life, Wellcome Sanger Institute, Hinxton, Cambridge, CB10 1SA, UK

18 Northeastern University, Boston, MA 02115, USA

19 University of Southern California, Quantitative and Computational Biology, Los Angeles, CA, USA

20 European Molecular Biology Laboratory, European Bioinformatics Institute, Wellcome Genome Campus, Cambridge, CB10 1SD, UK

21 Coriell Institute for Medical Research, Camden, NJ 08103, USA

22 Department of Computer Science, University of Pisa, Pisa 56127, Italy

23 Center for Health Data Science, University of Copenhagen, Denmark

24 Division of Medical Genetics, University of Washington School of Medicine, Seattle, WA 98195, USA

25 Department of Human Genetics, McGill University, Montreal, Québec H3A 0C7, Canada

26 Canadian Center for Computational Genomics, McGill University, Montreal, Québec H3A 0G1, Canada

27 Institute for the Advanced Study of Human Biology (WPI-ASHBi), Kyoto University, Kyoto 606-8501, Japan

28 Genome Informatics Section, Computational and Statistical Genomics Branch, National Human Genome Research Institute, National Institutes of Health, Bethesda, MD 20892, USA

29 Material Measurement Laboratory, National Institute of Standards and Technology, Gaithersburg, MD 20877, USA

30 Howard Hughes Medical Institute, Chevy Chase, MD 20815, USA

31 The Rockefeller University, New York, NY 10065, USA

32 Department of Genetics, Washington University School of Medicine, St. Louis, MO 63110, USA

33 Center for Genomic Health, Yale University School of Medicine, New Haven, CT 06510, USA

34 Google LLC, 1600 Amphitheater Pkwy, Mountain View, CA 94043, USA

35 School of Biological Sciences, Washington State University, Pullman WA 99163, USA

36 The Vertebrate Genome Laboratory, The Rockefeller University, New York, NY 10065, USA

37 National Institutes of Health (NIH)-National Human Genome Research Institute, Bethesda, MD, USA

38 Center for Computational and Genomic Medicine, The Children’s Hospital of Philadelphia, Philadelphia, PA 19104, USA.

39 Department of Biomolecular Engineering, University of California, Santa Cruz, 1156 High St., Santa Cruz, CA 95064, USA

40 Dovetail Genomics, Scotts Valley, CA 95066, USA

41 Division of Oncology, Department of Medicine, Stanford University School of Medicine, Stanford, CA, 94305, USA

42 European Molecular Biology Laboratory, Genome Biology Unit, Meyerhofstr. 1, 69117 Heidelberg, Germany

43 Computer Sciences Department, Barcelona Supercomputing Center, Barcelona, Spain

44 Departament d’Arquitectura de Computadors i Sistemes Operatius, Universitat Autònoma de Barcelona, Barcelona, Spain

45 Department of Biomedical Engineering, Johns Hopkins University, Baltimore 21218, MD, USA

46 Department of Ecology & Evolutionary Biology, University of California, Santa Cruz, 1156 High St, Santa Cruz, CA, USA

47 Berlin Institute for Medical Systems Biology, Max Delbrück Center for Molecular Medicine in the Helmholtz Association, Berlin, Germany

48 National Center for Biotechnology Information, National Library of Medicine, National Institutes of Health, Bethesda, MD 20894, USA

49 Al Jalila Genomics Center of Excellence, Al Jalila Children’s Specialty Hospital, Dubai, UAE

50 Center for Genomic Discovery, Mohammed Bin Rashid University of Medicine and Health Sciences, Dubai, UAE

51 Center for Computational Biology, Johns Hopkins University, Baltimore, MD 21218, USA

52 Department of Public Health Sciences, University of California, Davis, One Shields Avenue, Medical Sciences 1C, Davis, CA 95616

## Contributions

The study was conceived and designed by HJ.L and H.P.J. The k-mer indexing of all assemblies were generated by D.S.P. Data analysis was performed by HJ.L and S.U.G. The SV plots were generated by S.U.H, HJ.L and B.Z. The 1KGP SV overlap analysis was done by B.Z. and C.R.H. The manuscript was written by HJ.L. and H.P.J. HJ.L and H.P.J. managed the project.

## ETHICS DECLARATION

All authors declare no competing interests.

## Notes

https://dna-discovery.stanford.edu/publicmaterial/datasets/pangenome/

### Competing Interest Statement

The authors have declared no competing interest.

### Summary of Updates

The supplementary tables were added.

## REFERENCES

1 Sherman, R. M. & Salzberg, S. L. Pan-genomics in the human genome era. Nat Rev Genet 21, 243–254, doi:10.1038/s41576-020-0210-7 (2020).

2 Hurgobin, B. & Edwards, D. SNP Discovery Using a Pangenome: Has the Single Reference Approach Become Obsolete? Biology (Basel) 6, doi:10.3390/biology6010021 (2017).

3 Miga, K. H. & Wang, T. The Need for a Human Pangenome Reference Sequence. Annu Rev Genomics Hum Genet 22, 81–102, doi:10.1146/annurev-genom-120120-081921 (2021).

4 Nurk, S. et al. The complete sequence of a human genome. Science 376, 44–53, doi:10.1126/science.abj6987 (2022).

5 Wang, T. et al. The Human Pangenome Project: a global resource to map genomic diversity. Nature 604, 437–446, doi:10.1038/s41586-022-04601-8 (2022).

6 Liao, W.-W. et al. A Draft Human Pangenome Reference. bioRxiv, 2022.2007.2009.499321, doi:10.1101/2022.07.09.499321 (2022).

7 Kille, B., Balaji, A., Sedlazeck, F. J., Nute, M. & Treangen, T. J. Multiple genome alignment in the telomere-to-telomere assembly era. Genome Biol 23, 182, doi:10.1186/s13059-022-02735-6 (2022).

8 Pavlichin, D. S. et al. KmerKeys: a web resource for searching indexed genome assemblies and variants. Nucleic Acids Res, doi:10.1093/nar/gkac266 (2022).

9 Lau, B. T. et al. Profiling SARS-CoV-2 mutation fingerprints that range from the viral pangenome to individual infection quasispecies. Genome Med 13, 62, doi:10.1186/s13073-021-00882-2 (2021).

10 Lee, H., Shuaibi, A., Bell, J. M., Pavlichin, D. S. & Ji, H. P. Unique k-mer sequences for validating cancer-related substitution, insertion and deletion mutations. NAR Cancer 2, zcaa034, doi:10.1093/narcan/zcaa034 (2020).

11 Karczewski, K. J. et al. The mutational constraint spectrum quantified from variation in 141,456 humans. Nature 581, 434–443, doi:10.1038/s41586-020-2308-7 (2020).

12 Blanchette, M. et al. Aligning multiple genomic sequences with the threaded blockset aligner. Genome Res 14, 708–715, doi:10.1101/gr.1933104 (2004).

13 Kent, W. J. et al. The human genome browser at UCSC. Genome Res 12, 996–1006, doi:10.1101/gr.229102 (2002).

14 Morales, J. et al. A joint NCBI and EMBL-EBI transcript set for clinical genomics and research. Nature 604, 310–315, doi:10.1038/s41586-022-04558-8 (2022).

15 Abel, H. J. et al. Mapping and characterization of structural variation in 17,795 human genomes. Nature 583, 83–89, doi:10.1038/s41586-020-2371-0 (2020).

16 Byrska-Bishop, M. et al. High-coverage whole-genome sequencing of the expanded 1000 Genomes Project cohort including 602 trios. Cell 185, 3426–3440 e3419, doi:10.1016/j.cell.2022.08.004 (2022).

17 Genomes Project, C. et al. A global reference for human genetic variation. Nature 526, 68–74, doi:10.1038/nature15393 (2015).

18 Delage, W. J., Thevenon, J. & Lemaitre, C. Towards a better understanding of the low recall of insertion variants with short-read based variant callers. BMC Genomics 21, 762, doi:10.1186/s12864-020-07125-5 (2020).

19 Mahmoud, M. et al. Structural variant calling: the long and the short of it. Genome Biol 20, 246, doi:10.1186/s13059-019-1828-7 (2019).

20 Wagner, J. et al. Curated variation benchmarks for challenging medically relevant autosomal genes. Nat Biotechnol 40, 672–680, doi:10.1038/s41587-021-01158-1 (2022).

