## Supplementary Figures for "The Human Pangenome’s sequence conservation reveals a landscape of polymorphic structural variations"

**SUPPLEMENTARY NOTES**

**1. The pan-conserved segment tags (PSTs) in all assemblies**

The total number of PSTs used in this study was ~13.48 x $10^6$ and checked if the sequences of them were maintained in same way across all assemblies. To confirm the preservation of PSTs across all assemblies, we evaluated the lengths between following number of non-overlapping 31-mers within pan-conserved segments: ***ceil(L/31)*** where ceil is the ceiling function and L is the length of pan-conserved segments. For instance, we checked 3 of non-overlapping constituent 31-mers of a PST with size of 100 bp. We found out that the lengths of 237 PSTs were different in several assemblies (**Supplementary Table S11)**. This happened when one of constituent pan-conserved 31-mer tag were accidently identical to a 31-mer from other location due to SNP.

**2. The landscape of divergent lengths on assemblies relative to GRCh38**

In overall, we observed the number of longer divergent lengths (indicating insertion) is larger than shorter divergent lengths (indicating deletion) **(Supplementary Table S12)**. For interval lengths ranging up to 1 Kb, we observed minimal divergence in the length, meaning that the same size intervals were observed across all haploid genomes. As expected, longer interval lengths among the Pangenome assemblies diverged more frequently from GRCh38. Interestingly, intervals in the size range of 1Kb to 10 Kb bracket or the 10 Mb to 100 Mb brackets were generally shorter than what was calculated from GRCh38.

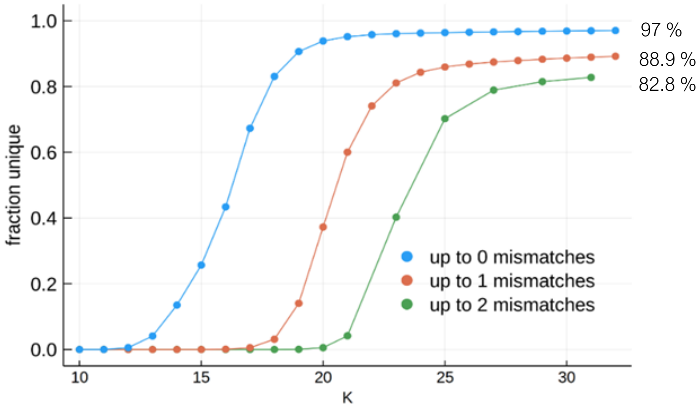


**Supplementary Figure 1**. The extent of uniqueness by k-mer length within 2 mismatches.

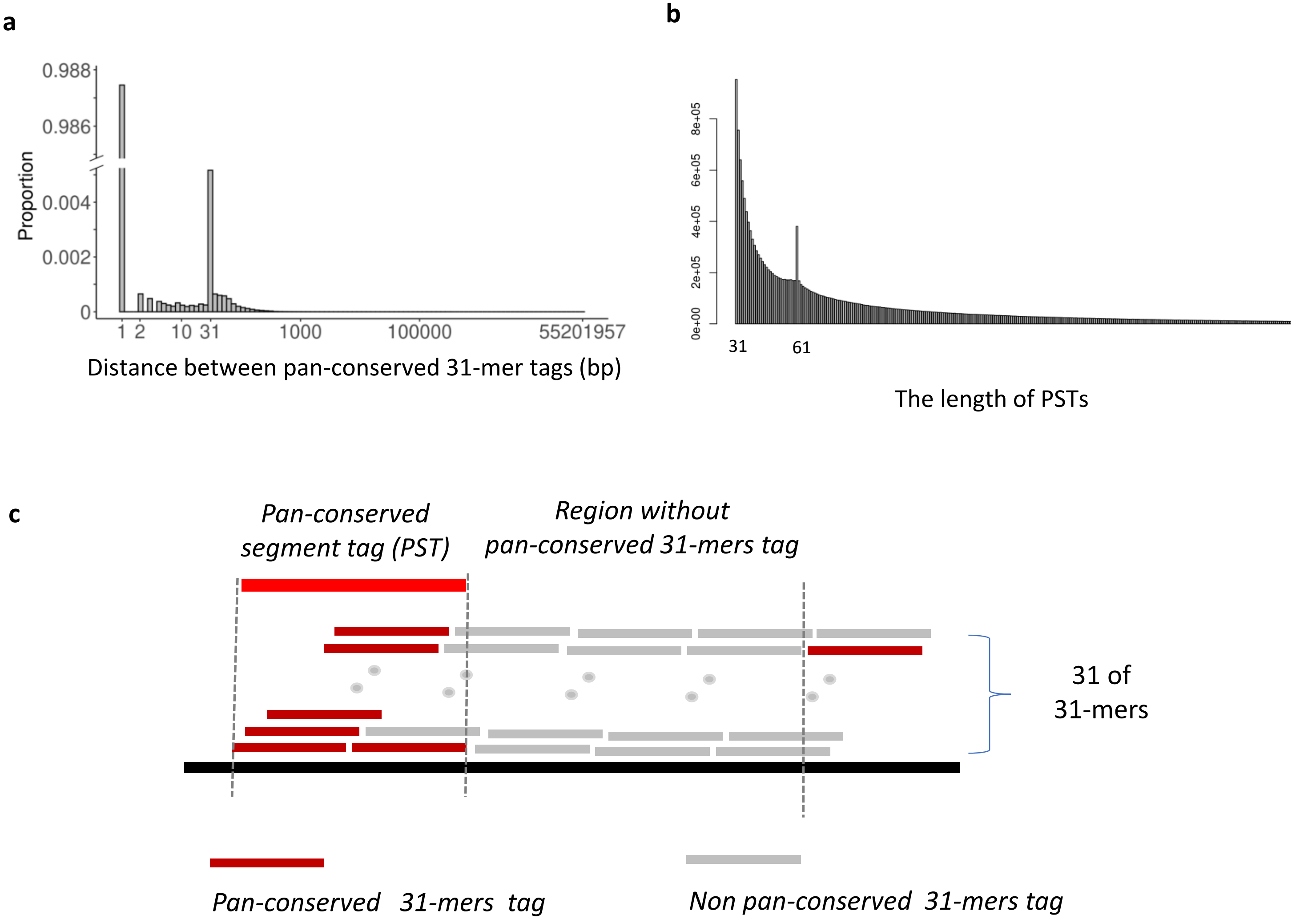


**Supplementary Figure 2**. The spatial distribution of pan-conserved 31-mers based on GRCH38 coordinates. **a**, The proportion of distance between adjacent 31-mer pairs. Most of pan-conserved 31-mer are consecutive and another peak at the distance of 31, which could be explained by the presence of single nucleotide polymorphism (**SNP**) across assemblies. **b**, The distribution of the length of pan-conserved segments. **c**, "**pan-conserved segment tag**", stretch of consecutive pan-conserved 31-mer tags, and regions with depletion of pan-conserved 31-mer tags

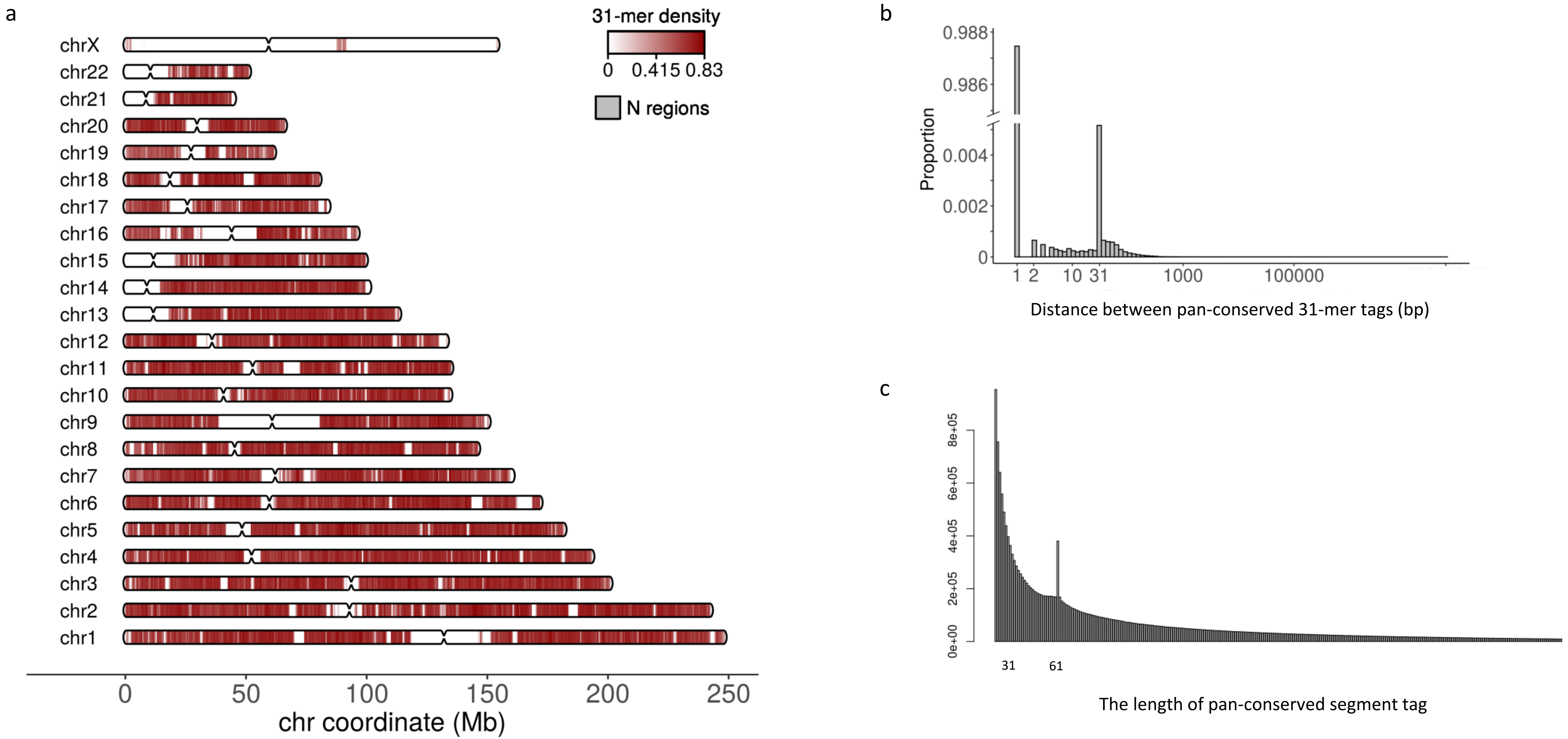


**Supplementary Figure 3**. a, The distribution of pan-conserved segment tags (PSTs) on CHM13. b, The proportion of distance between adjacent 31-mer pairs. c, The distribution of the length of pan-conserved segments

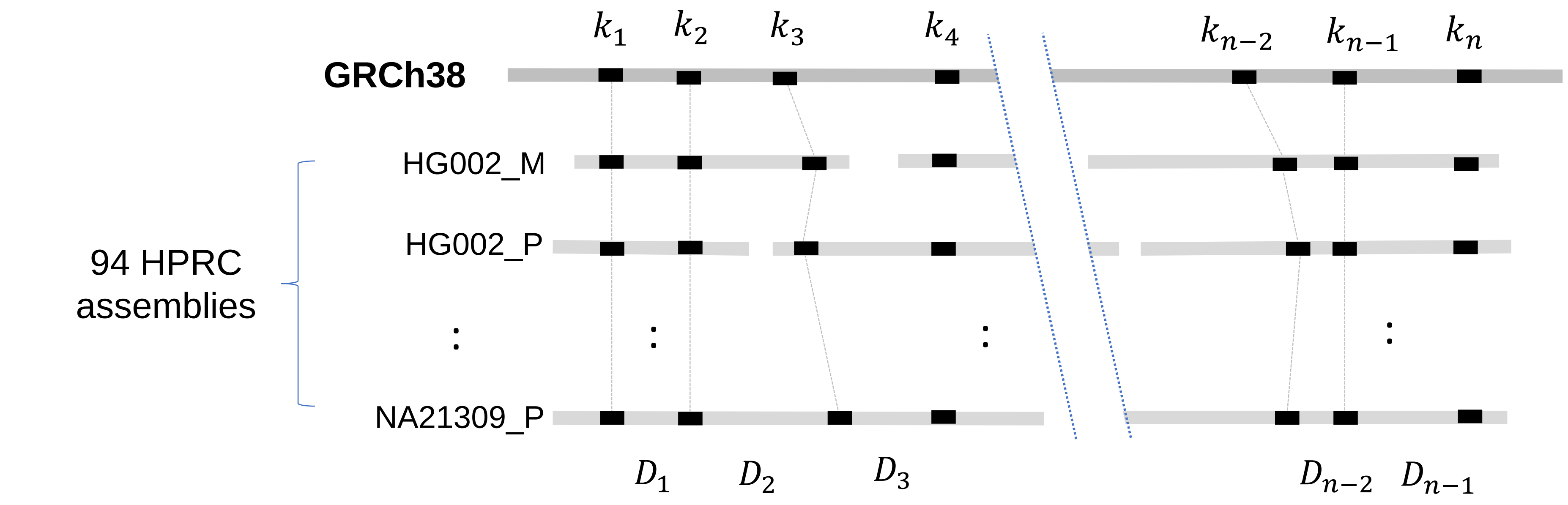


| | | $D_1$ | $D_2$ | $D_3$ | - - | $D_{n-2}$ | $D_{n-1}$ |
|---|---|---|---|---|---|---|---|
| | ***GRCh38*** | ***70*** | ***332*** | ***530*** | - - | ***400*** | ***450*** |
| Distance matrix | HG002 maternal | 70 | 460 | NA | - - | 310 | 450 |
| | HG002 paternal | 70 | 445 | 500 | - - | 200 | 450 |
| | : | : | : | : | : | : | : |
| | NA21309 paternal | 70 | 600 | 460 | - - | 260 | 450 |
| Summary | Num. same distances | 94 | 82 | 30 | - - | 0 | 94 |
| | Num. different distance | 0 | 11 | 62 | - - | 94 | 0 |
| | Median differences | 0 | 152 | -80 | - - | -200 | 0 |

Constant Intervals    Polymorphic Intervals

**Supplementary Fig 4**. The matrix for interval lengths between pan-conserved sequences across all 94 HPRC assemblies. The distance between tandem pan-conserved sequences cannot be measured when they are on different contigs, indicated as 3000001 (meaning NA). The transpose of this matrix is provided as supplementary file 2.

**Supplementary Fig 5**

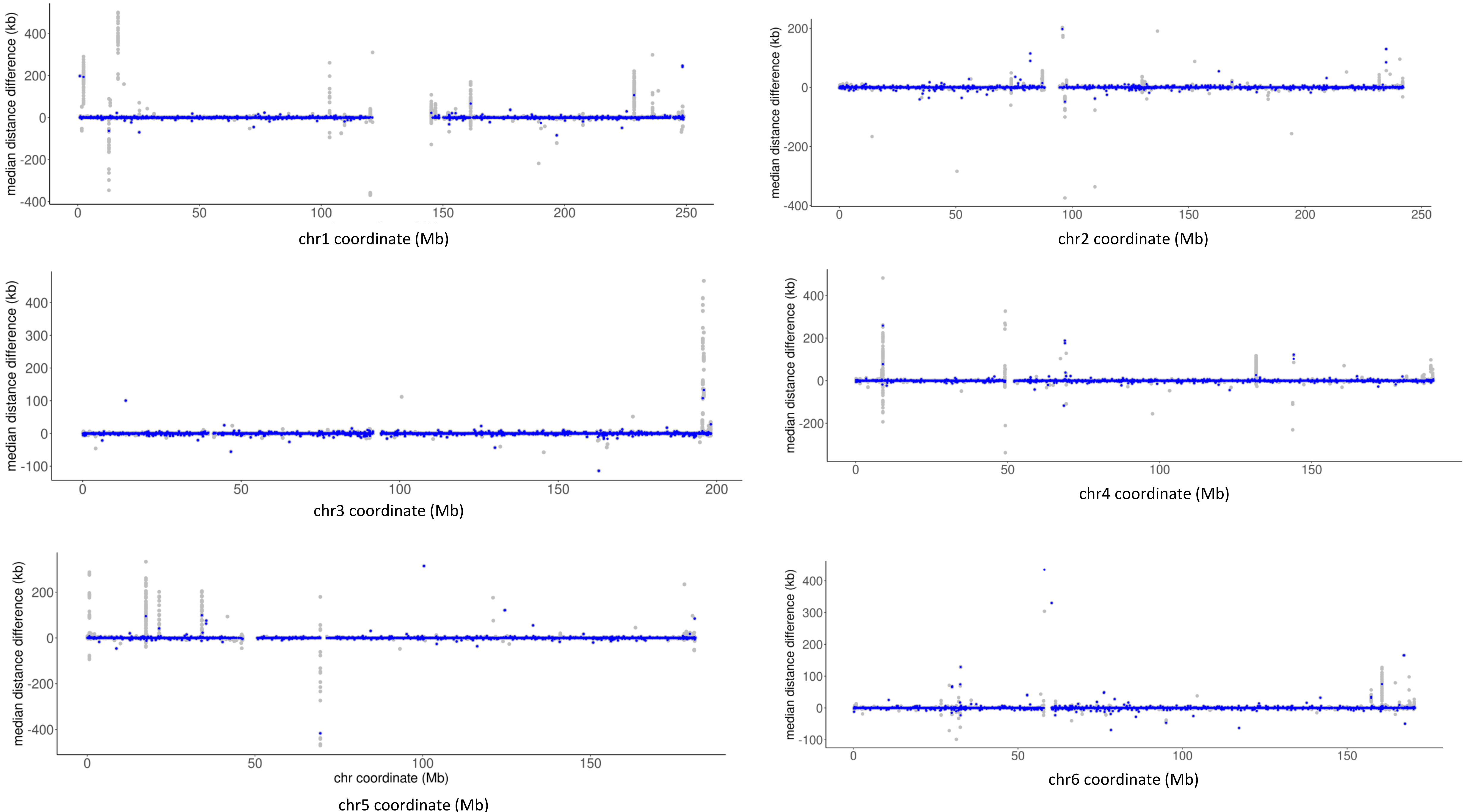

**Supplementary Fig 5**

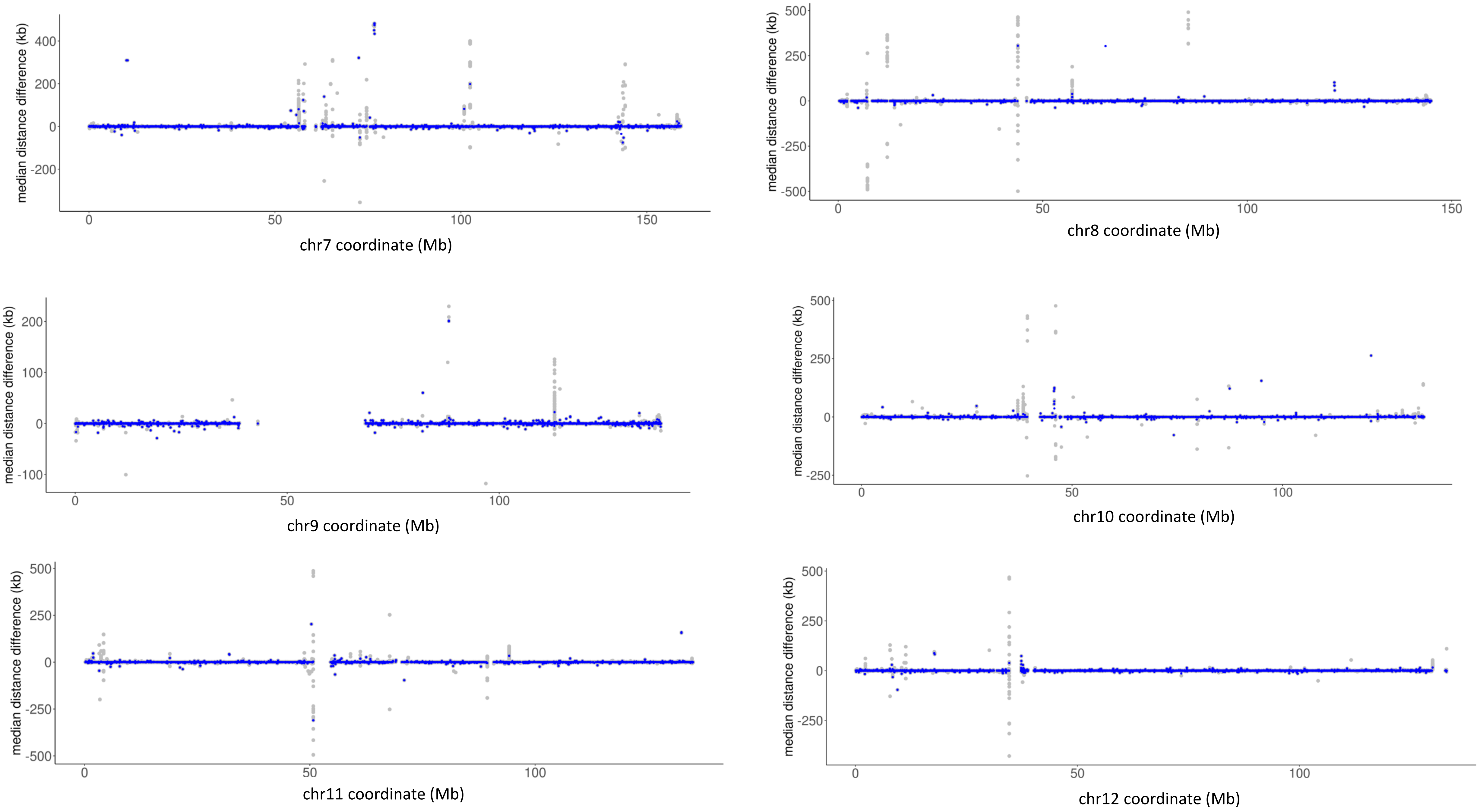

**Supplementary Fig 5**

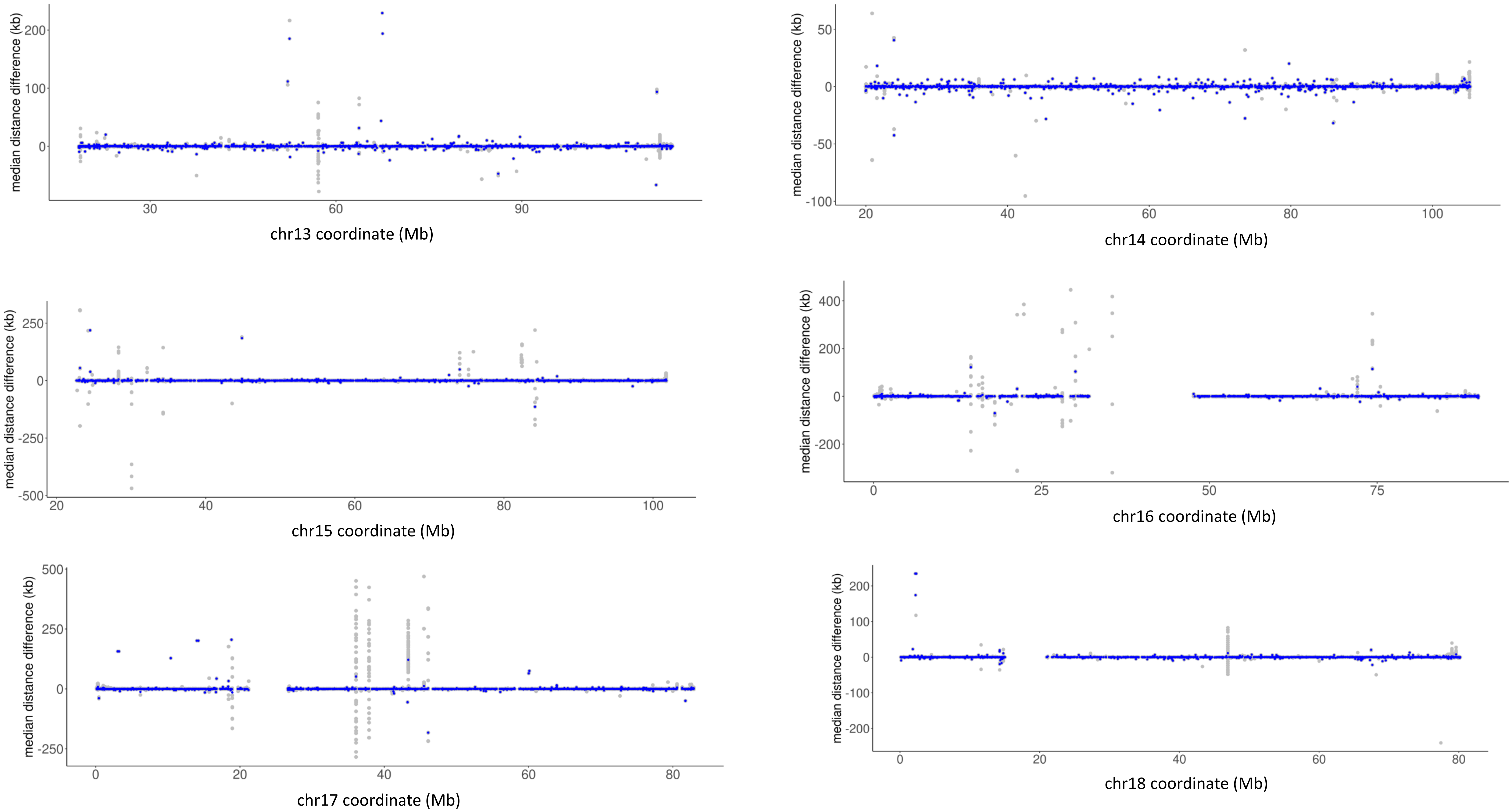

**Supplementary Fig 5**

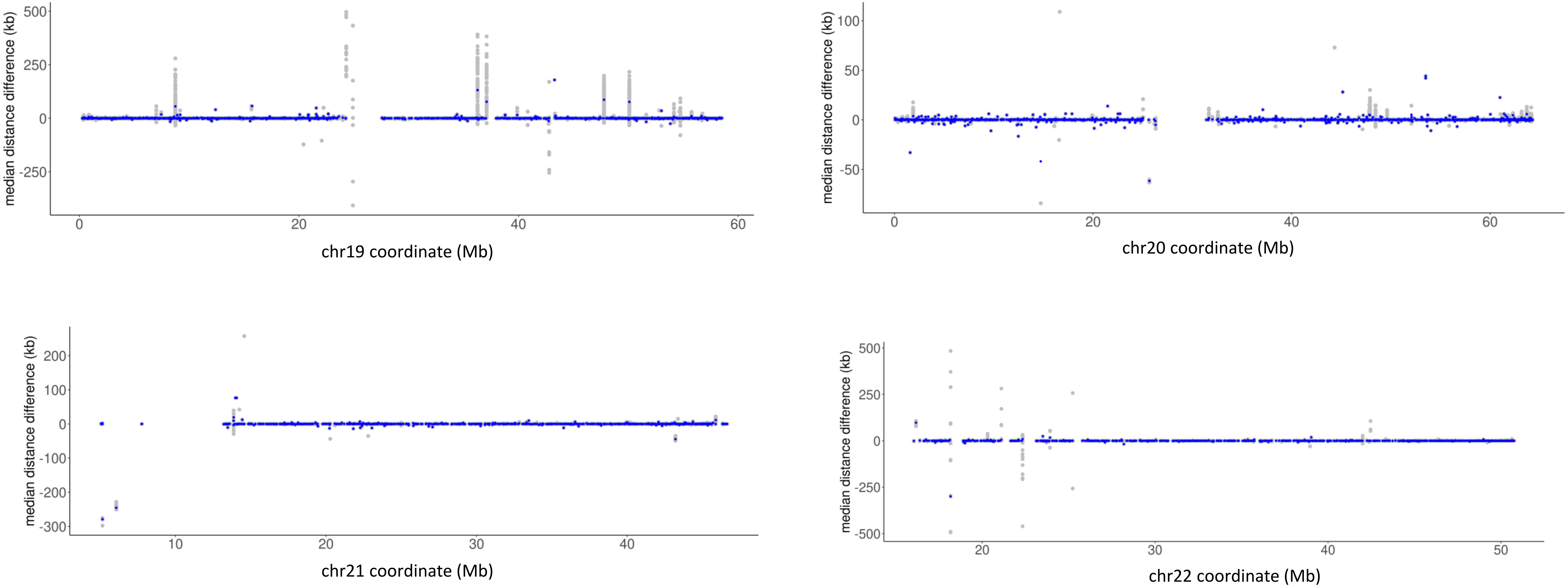

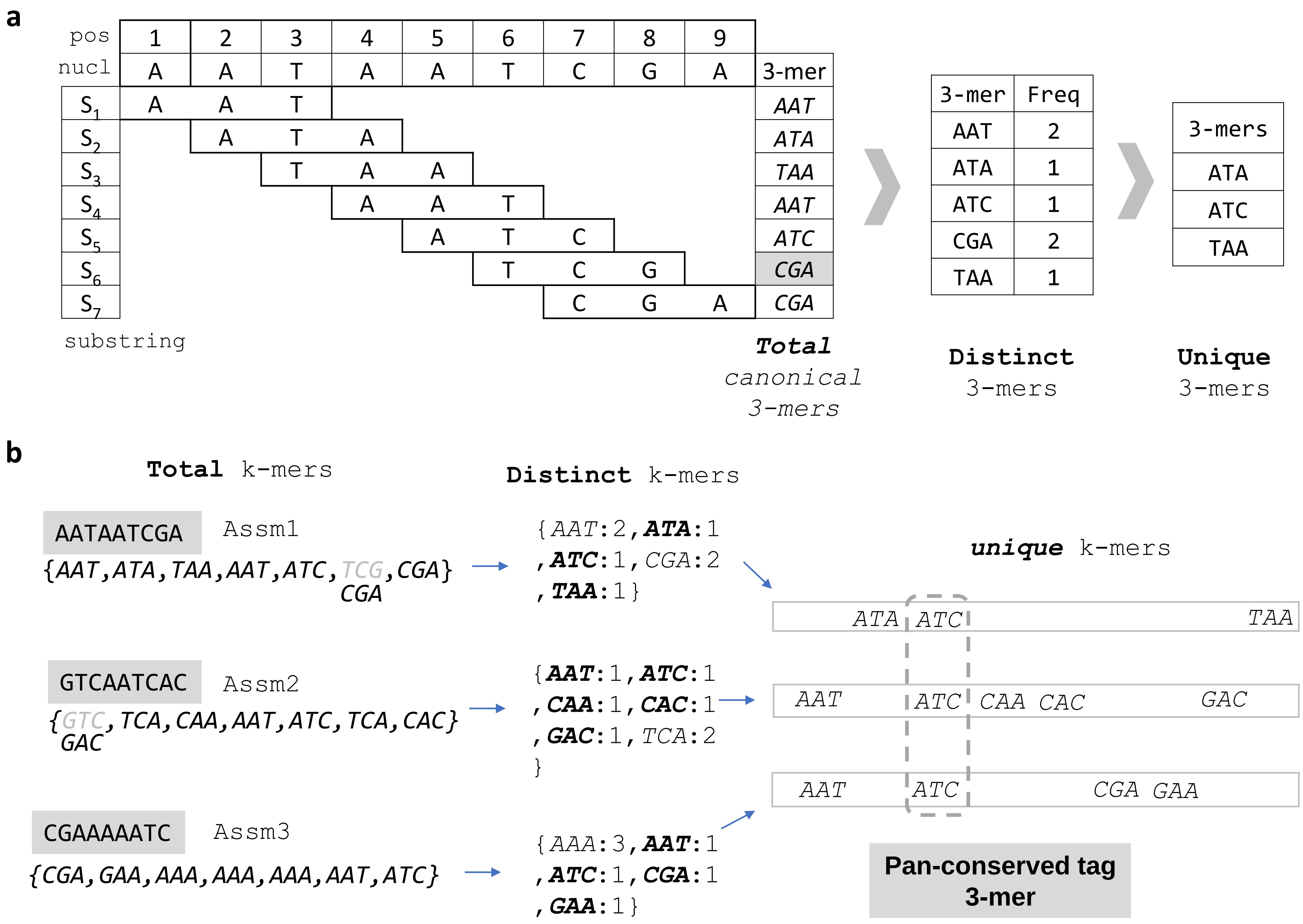


**Supplementary Fig. 6**. The indexing of assembly using k-mers. a, Different type of k-mers, b, The identification of pan-conserved sequence tag (PST)
